# RNAlater and flash freezing storage methods nonrandomly influence observed gene expression in RNAseq experiments

**DOI:** 10.1101/379834

**Authors:** Courtney N. Passow, Thomas J. Y. Kono, Bethany A. Stahl, James B. Jaggard, Alex C. Keene, Suzanne E. McGaugh

## Abstract

RNA-sequencing is a popular next-generation sequencing technique for assaying genome-wide gene expression profiles. Nonetheless, it is susceptible to biases that are introduced by sample handling prior gene expression measurements. Two of the most common methods for preserving samples in both field-based and laboratory conditions are submersion in RNAlater and flash freezing in liquid nitrogen. Flash freezing in liquid nitrogen can be impractical, particularly for field collections. RNAlater is a solution for stabilizing tissue for longer-term storage as it rapidly permeates tissue to protect cellular RNA. In this study, we assessed genome-wide expression patterns in 30 day old fry collected from the same brood at the same time point that were flash-frozen in liquid nitrogen and stored at −80°C or submerged and stored in RNAlater at room temperature, simulating conditions of fieldwork. We show that sample storage is a significant factor influencing observed differential gene expression. In particular, genes with elevated GC content exhibit higher observed expression levels in liquid nitrogen flash-freezing relative to RNAlater-storage. Further, genes with higher expression in RNAlater relative to liquid nitrogen experience disproportionate enrichment for functional categories, many of which are involved in RNA processing. This suggests that RNAlater may elicit a physiological response that has the potential to bias biological interpretations of expression studies. The biases introduced to observed gene expression arising from mimicking many field-based studies are substantial and should not be ignored.

## Introduction

High throughput sequencing technologies, such as RNA-sequencing methods, have revolutionized the quantification of genome-wide expression patterns across a broad range of fields in biological sciences (López-Maury *et al.* 2008; Wang *et al.* 2009). However, storage and RNA extraction methods prior to RNA-seq library preparation exert substantial impacts on biological studies, and often account for the majority of variation in a dataset if conditions and protocols are not identical across all samples (Todd *et al.* 2016). With the rise of RNAlater (Ambion, Invitrogen) as a popular storage method in field-based studies (De Smet *et al.* 2017; Wille *et al.* 2018), it is important to quantify if there are systematic biases in gene expression when samples are preserved in RNAlater versus flash-frozen in liquid nitrogen. In our literature review, however, we could find few direct comparisons of RNAseq data obtained from the most common field-preservation method RNAlater and the “gold standard” of flash freezing samples in liquid nitrogen (Alvarez *et al.* 2015; Wolf 2013) (but see(Cheviron *et al.* 2011; Choi *et al.* 2016)). Further, no studies examined whether a systematic bias due to gene characteristics exists for samples preserved in RNAlater.

Currently, two of the most common methods for RNA preservation and storage are flash freezing in liquid nitrogen and preservation in aqueous sulfate salt solutions, such as commercially available RNAlater. Flash freezing, usually through the use of immersing the sample in dry ice or liquid nitrogen, is the most preferred means of stabilizing tissue samples for downstream analysis (Wolf 2013). While preferred, it can often be difficult to access and transport dry ice or liquid nitrogen, particularly in field conditions (Mutter *et al.* 2004). Hence, in the past decade, it has become common practice, especially in field environments, to store RNAseq-destined samples in RNAlater, a stabilizing solution that minimizes the need to readily process samples or chill the tissue. RNAlater can rapidly permeate tissue to stabilize and protect RNA (Chowdary *et al.* 2006; Florell *et al.* 2001). Likewise, RNAlater-immersed samples can be stored safely at room temperature for a week, and longer when stored at colder temperatures. Though, common practice is to store samples in RNAlater in field conditions for much longer than a week. While the exact ingredients of commercial RNAlater are unknown, the Material Safety Data Sheet lists inorganic salt as the major component and the homemade versions contain ammonium sulfate, sodium citrate, ethylenediaminetetraacetic acid (EDTA), and adjustment of pH using sulfuric acid.

In this study, we quantified the effects of storage condition on gene expression and examined differentially expressed genes for specific characteristics to assay for systematic bias. Individual, Mexican tetra fry (*Astyanax mexicanus*), were collected from the same brood and stored immediately in liquid nitrogen (N = 6) or RNAlater (N = 5). We specifically asked (1) Does storage condition affect patterns of differential gene expression and if so, (2) Are these effects on gene expression non-random, such that genes with certain features are differentially affected by storage condition? We found that a majority of the variation in gene expression was explained by storage condition. Likewise, we found that genes with higher GC content exhibited higher expression values in liquid nitrogen than RNAlater. Based on these findings, RNAlater-storage may potentially bias biological conclusions of RNAseq experiments.

## Methods

### Sample Collection

Samples for the transcriptome analyses were collected from a surface population of *Astyanax mexicanus* (total of 8 parents) that had been reared in the Keene laboratory at Florida Atlantic University for multiple generations. Parental fish were derived from wild-caught Río Choy stocks originally collected by William Jeffery. To minimize variation outside of storage methods, all individuals were collected from the same clutch (fertilized on 2016-12-08). Fish were raised in standard conditions, three days prior to experiment, fish were transferred into dishes with 12-21 fish per dish in a 14:10 light-dark cycle. Individuals were raised for 30 days after fertilization under standard conditions, when five individuals were sampled and stored in RNAlater and six individuals were flash frozen in liquid nitrogen and stored at −80. These fish were a part of a larger experiment and so for 24 hours prior to sampling, fish were kept in total darkness and sampled at 16:00h (10pm). To mimic field conditions, RNAlater individuals were stored at room temperature for 17 days (Camacho-Sanchez *et al.* 2013; Kono *et al.* 2016). Procedures for all experiments performed were approved by the Institutional Animal Care and Use Committee at Florida Atlantic University (Protocol #A15-32).

### RNA extraction, library preparation and sequencing

For RNA isolation, all individuals were processed within a week of each other (between 2017-01-19 and 2017-01-24), and RNAlater stored individuals were processed 17 days after initial storage (2017-01-24) (Table S1) with the same researcher performing all extractions. Whole organisms (< 30 mg of tissue) were homogenized using Fisherbrand pellet pestles and cordless motor (Fisher Scientific) in the lysate buffer RLT plus. Total RNA was extracted using the Qiagen RNAeasy Plus Mini Kit (Qiagen) and quantified using NanoDrop Spectrophotometer (Thermo Fisher Scientific), Ribogreen (Thermo Fisher Scientific), and Bioanalyzer (Agilent) to obtain RNA integrity numbers (RIN). All cDNA libraries were constructed at the University of Minnesota Genomics Center on the same day in the same batch. In brief, a total of 400 ng of RNA was used to isolated mRNA via oligo-dT purification. dsDNA was constructed from the mRNA by random-primed reverse transcription and second-strand cDNA synthesis. Strand-specific cDNA libraries were then constructed using TruSeq Nano Stranded RNA kit (Illumina), following manufacturer protocol. Library quality was assessed using Agilent DNA 1000 kit on a Bioanalyzer (Agilent). To minimize batch effects, barcoded libraries were then pooled and sequenced across multiple lanes of an Illumina HiSeq 2500 to produce 125-bp paired-end reads at University of Minnesota Genomics Center (Table S1). All sequence data were deposited in the short read archive (Study Accession ID: RNAlater: SRX3446133, SRX3446136, SRX3446135, SRX3446155, SRX3446156; liquid N2: SRS2736519, SRS2736520, SRS2736523, SRS2736524, SRS2736525,SRS2736526).

### RNAseq quality check

The raw RNA-seq reads were quality checked using Fastqc (Andrews 2014) and trimmed to removed adapters using the program Trimmomatic version 0.33; (Bolger *et al.* 2014). Trimmed reads were mapped to the *Astyanax mexicanus* reference genome (version 1.0.2; GenBank Accession Number: GCA_000372685.1; (McGaugh *et al.* 2014). Mapping was conducted using the splice-aware mapper STAR (Dobin *et al.*2013), because it yielded the higher alignment percentage and quality compared to a similar mapping program (HISAT2, results not shown (Kim *et al.* 2015)). We used Stringtie (version 1.3.3d; (Pertea *et al.* 2015) (Pertea *et al.* 2016) to quantify number of reads mapped to each gene in the reference annotation set of the *A. mexicanus* genome, and used the python script provided with Stringtie (prepDE.py) to generate a gene counts matrix (Pertea *et al.* 2016). R (Team 2014) was used to compare RIN between liquid nitrogen and RNAlater treatments using a nonparametric Kruskal-Wallis test.

### Variation in gene expression

To visualize changes in observed gene expression, we performed principal components analysis on a gene counts matrix. Genes with less than 100 counts across all samples were removed from the matrix because genes with low counts bias the differential expression tests (Love *et al.* 2014). The resulting counts were decomposed into a reduced dimensionality data set with the prcomp() function in R (Team 2014).

To identify genes that showed the largest difference in observed gene expression between storage conditions, we performed a differential expression analysis between samples flash frozen in liquid nitrogen (N = 6) and samples stored in RNAlater (N = 5) using DESeq2 (Love *et al.* 2014). DESeq2 normalizes expression counts for each sample and then fits a negative binomial model for counts for each gene. Samples with the same storage condition were treated as replicates, (i.e., the variation due to storage was assumed to be greater than variation among biological samples). This was confirmed in the PCA plot (Figure 1), where PC1 linearly separated samples based on their treatments. P-values for differential expression were adjusted based on the Benjamini-Hochberg algorithm, using a default false discovery rate of at most 0.1 (Love *et al.* 2014). Genes were labeled as differentially expressed if the Benjamini-Hochberg adjusted P-value was less than 0.1. Log2(RNAlater/liquid nitrogen) values were calculated with DESeq2, and exported for further analysis.

**Figure 1:**
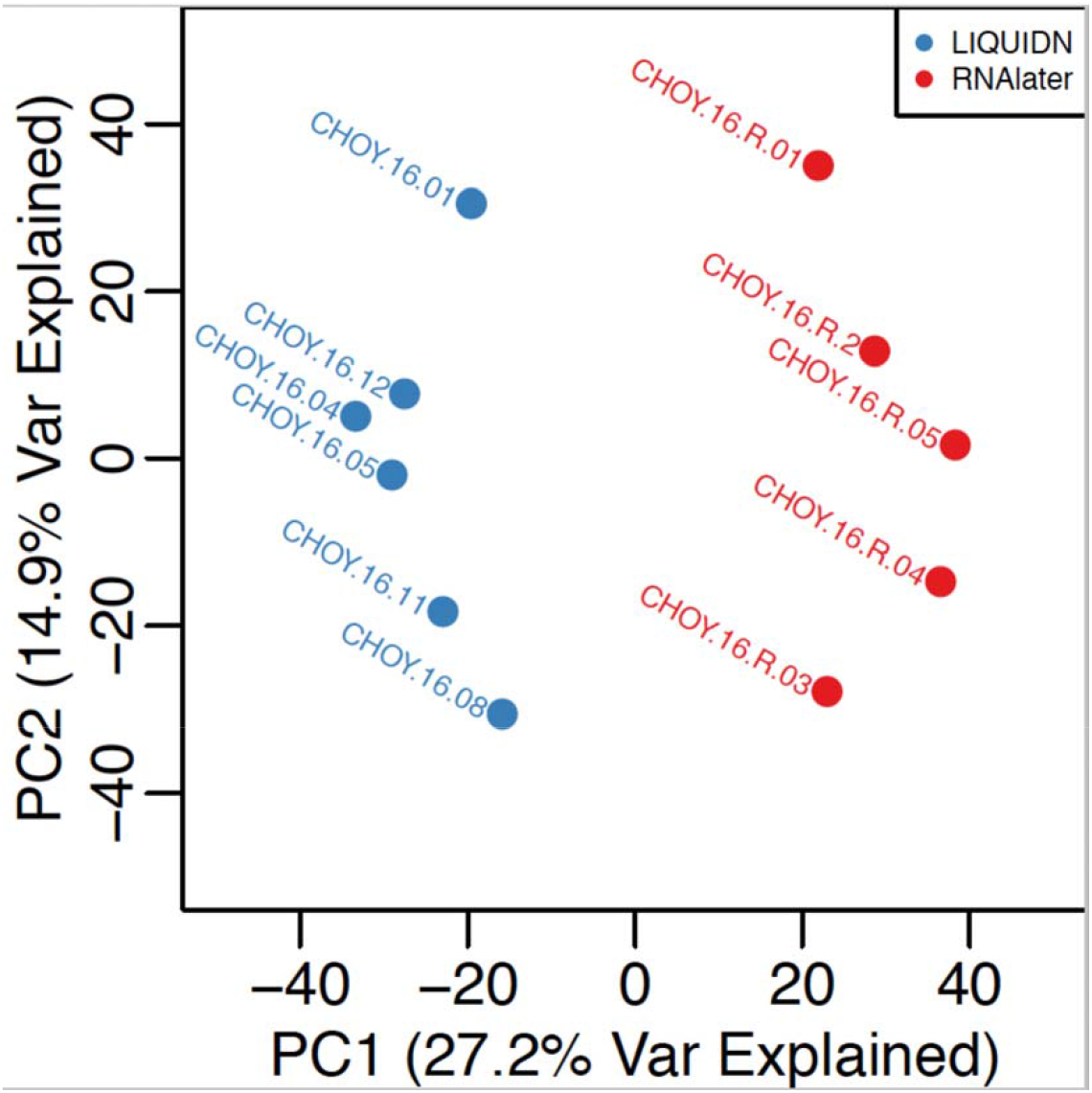
Principal components analysis plot showing PC1 and PC2 for each sample. RNAlater samples (red) are linearly separated from liquid nitrogen samples (blue) by PC1.

Linear model to determine factors influencing differential expression To identify the factors that contribute to the variability in gene expression between preservation methods, we fit a linear model of observed gene expression of all genes as a function of various genomic characteristics. We tested the contributions of mean expression level, annotated gene length, exon number, GC content, presence or absence of simple sequence repeats, and presence or absence of a homopolymer tract to differences in observed gene expression between preservation methods. We used the log2(RNAlater/liquid nitrogen) values from DESeq2 as the measure of change in observed gene expression, and the mean of normalized counts as the mean expression level. The annotated gene length was calculated as the total length of the gene annotation, including noncoding (i.e., intronic) regions. A simple sequence repeat was defined as two or more nucleotides repeated at least three times in tandem, and a homopolymer tract was defined as a single nucleotide repeated at least six times in tandem in the reference genome. Repeat presence or absence was based only on the reference genome sequence, and were not scored to be polymorphic in the sample. Length and exon number were calculated with a modified version of GTFTools (Li 2018). GC content, presence/absence of a simple sequence repeat, and presence/absence of a homopolymer repeat were scored with custom Python scripts available on our GitHub repository. Notably, the reference genome was based off the Pachón cavefish, and it is conceivable that some homopolymers and sequence repeats may not be identical in the surface fish.

We performed model selection on a series of linear models using likelihood ratio tests of nested models. The “full model” was as follows:

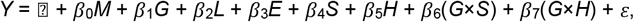

where *Y* is log2(RNAlater/liquid nitrogen) of expression between treatments, *M* is the the normalized mean expression value across all samples, *G* is GC content, *L* is gene length, *E* is the total number of exons in the gene, S is SSR presence/absence, and *H* is homopolymer presence/absence. GC content, gene length, and exon number were treated as continuous variables, and SSR presence and homopolymer presence were treated as categorical variables. Model selection proceeded by testing the contributions of the interaction terms to the variance explained, and removing them if not significant. We tested the terms with the lowest non-significant t-values in the regression, and removed them if they did not significantly improve model fit.

### Annotation of differentially expressed genes

Since most of the variation was explained by a technical variable (i.e., preservation and storage), we did not expect biologically meaningful annotation. However, we conducted annotation analyses using two different methods. Differentially expressed genes at the 0.05 false discovery rate were converted to homologous zebrafish (*Danio rerio*) gene IDs for a gene ontology (GO) term enrichment analysis. Duplicate zebrafish gene IDs were removed prior to GO term enrichment analyses. GO term enrichment was tested with the GOrilla webserver (Eden *et al.* 2009) (http://cbl-gorilla.cs.technion.ac.il/), with a database current as of 2018-07-07. Other running parameters were left at their default values. In addition, PANTHER analysis (Mi *et al.* 2016) (http://pantherdb.org/tools/compareToRefList.jsp) was run using 1:1 orthologs between zebrafish and *Asytanax* with database current as of 2018-04-30. Within the PANTHER suite, we used PANTHER v13.1 overrepresentation tests (i.e., Fisher’s exact tests with FDR multiple test correction) with the Reactome v58, PANTHER proteins, GoSLIM, GO, and PANTHER Pathways. For both annotation analyses, they were run with two lists of unranked gene IDs: the target list was the differentially expressed gene IDs (either higher or lower expression in RNAlater), and the background list was all zebrafish genes genome-wide.

### Script Availability

Scripts to perform all data QC and processing are available at https://github.com/TomJKono/CaveFish_RNAlater

## Results

### Mapping statistics and annotation

RNA sequencing from whole, 30-days post fertilization individuals yielded a total of 108,874,500 reads for individuals stored in liquid nitrogen (mean = 18,145,750 ± stdev 1,938,410 per individual; N = 6) and 82,448,455 reads for individuals stored in RNAlater (mean = 16,489,691 ± stdev 1,890,519 per individual; N = 5) (Table 1). While all RIN scores passed the threshold (> 7), RIN scores were significantly different between RNAlater and liquid nitrogen treatments (Kruskal-Wallis chi-squared = 7.6744, df = 1, p-value = 0.005601; RNAlater mean RIN = 8.60, liquid nitrogen mean RIN = 9.83).

**Table 1:** Reported are the number of reads (after adapter trimming) used as input for the mapping software (STAR), number of reads that uniquely mapped to the reference genome, and the percent of reads that mapped to the reference genome.

| Sample Name | Treatment | Input reads | Uniquely mapped reads | % Mapped |
| --- | --- | --- | --- | --- |
| CHOY-16-01 | Liquid N2 | 20,162,412 | 18,125,738 | 89.90% |
| CHOY-16-04 | Liquid N2 | 15,760,631 | 13,812,190 | 87.64% |
| CHOY-16-05 | Liquid N2 | 18,025,208 | 16,015,383 | 88.85% |
| CHOY-16-08 | Liquid N2 | 16,368,007 | 14,584,314 | 89.10% |
| CHOY-16-11 | Liquid N2 | 17,997,036 | 15,126,300 | 89.61% |
| CHOY-16-12 | Liquid N2 | 20,561,206 | 18,221,558 | 88.62% |
| CHOY-16-R-01 | RNAlater | 17,984,846 | 15,643,479 | 86.98% |
| CHOY-16-R-03 | RNAlater | 17,064,911 | 14,913,653 | 87.39% |
| CHOY-16-R-04 | RNAlater | 13,585,649 | 11,809,525 | 86.93% |
| CHOY-16-R-05 | RNAlater | 15,692,250 | 13,716,160 | 87.41% |
| CHOY-16-R-2 | RNAlater | 18,120,799 | 15,851,038 | 87.47% |

Total yield of reads and number of uniquely mapping reads were not significantly different between treatments (*t* = 1.4301; *P* = 0.1875). Samples on average mapped 88.17% of the reads to the *Astyanax mexicanus* genome (range: 86.93%-89.90%), with liquid nitrogen samples mapping on average 88.17% and RNAlater mapping 87.24%.

Filtering of the gene counts matrix to include only genes with ≥100 reads resulted in 15,515 genes being used for both clustering and differential expression analysis. Annotations were extracted from the *Astyanax mexicanus* annotation file (Astyanax_mexicanus.AstMex102.91.gtf). Distributions of raw and filtered gene expression counts are given in Figure S1.

### PCA and Differentially Expressed Genes

Principal components analysis showed that the major axis of differentiation among the samples was treatment (Figure 1). This corresponds to the first principal component, and explains 27.2% of the variation. Beyond the first principal component, the samples do not cluster into further discernable sub-groups, suggesting that the main axis of differentiation among these samples is their storage conditions (Figure S2 A and B).

A total of 2,708 (17.5%) genes were significantly differentially expressed between treatments at the 0.05 significance level (Figure 2). Of these, 1,635 exhibited significantly lower observed expression in RNAlater than liquid nitrogen, and 1,073 exhibited significantly higher observed expression in RNAlater than liquid nitrogen.

**Figure 2:**
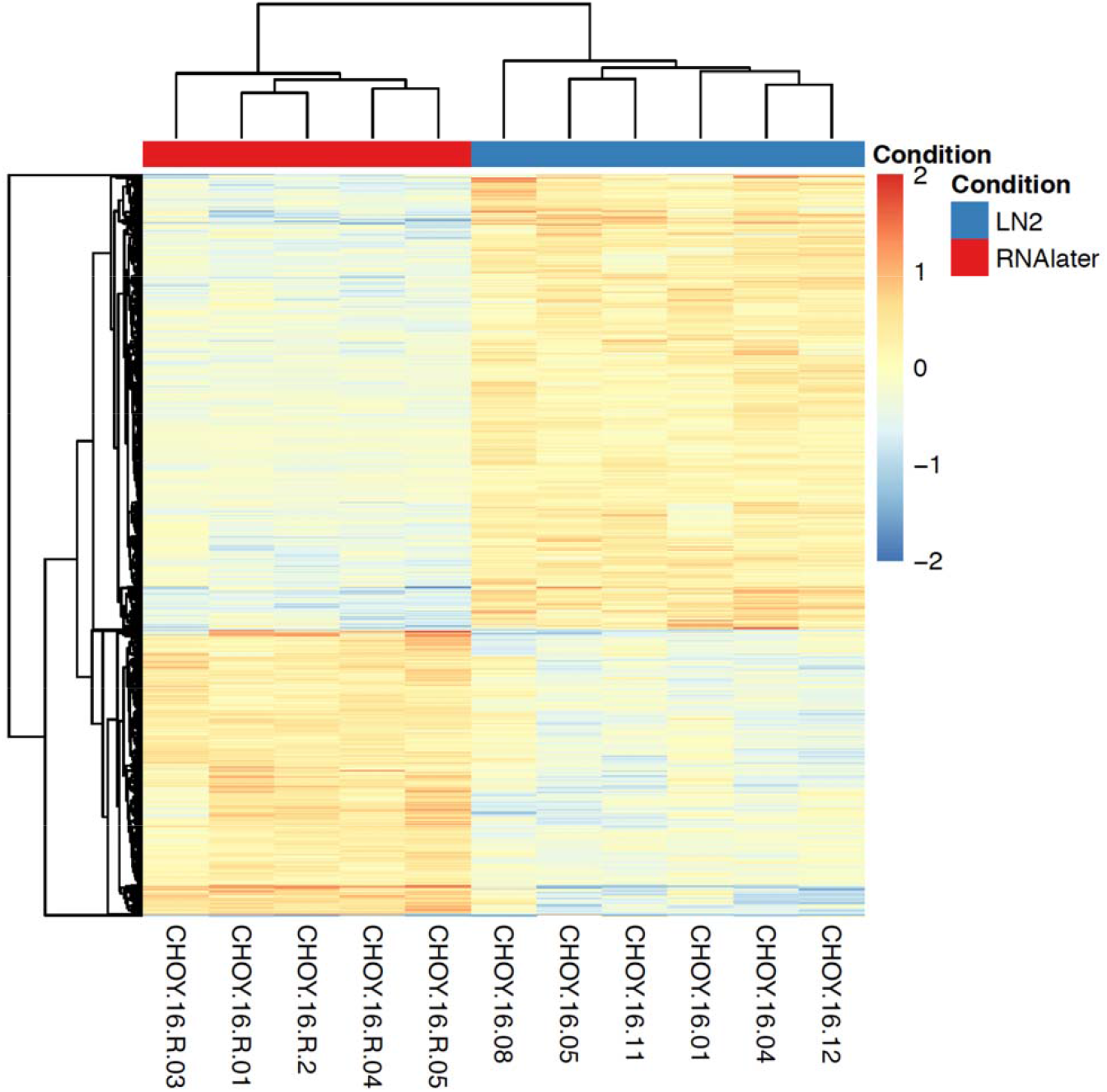
Clustering heatmap showing genes that are differentially expressed among RNAlater samples and liquid nitrogen samples. Gene expression values have been normalized by sample, then centred about 0 for each gene. This heatmap contains differentially expressed genes (after FDR correct with p < 0.05) including 1,073 genes that with higher expression values in the RNAlater treatment relative to the liquid nitrogen treatment, and 1,635 genes that exhibited lower expression values the RNAlater treatment.

### Annotation of differentially expressed genes

We expected little GO term enrichment as differences in gene expression would likely be due to differences in preservation techniques, not biological variation. Further, the number of enrichment categories for higher- and lower-expressed genes in RNAlater with respect to liquid nitrogen was similar across annotation programs. However, we observed substantially different functional enrichment among genes that were higher- and lower-expressed in RNAlater compared to liquid nitrogen across annotation programs.

In the GOrilla analyses, GO term enrichment analysis showed that the genes that were differentially expressed between treatments were spread across a broad range of GO terms. In genes that are significantly lower in RNAlater in comparison to liquid nitrogen, the only significantly enriched GO term is protein autophosphorylation (GO:0046777). In genes that are significantly higher in RNAlater, there were 13 enriched GO terms (after FDR correction; Supplementary Material). These included acyl-CoA, thioester, and sulfur compound metabolic processes, and purine nucleoside, nucleoside, and ribonucleoside bisphosphate metabolic processes. Notable, many of these processes involve replacing the linking oxygen in an ester by a sulfur atom, and if the homemade version of RNAlater is consistent with the commercial recipe, ammonium sulfate is likely the largest component.

The PANTHER suite annotation results were similar to the GOrilla analyses (Supplemental Materials). For genes that were significantly lower in RNAlater compared to liquid nitrogen, very few functional categories were enriched. However, many categories were significantly enriched for genes that were more highly expressed in RNAlater than liquid nitrogen. The most enriched categories in reactome pathways are involved in gene expression and processing of mRNA. Likewise, enriched PANTHER protein classes include RNA binding proteins, mRNA processing and splicing factors, and transcription factors. Enriched GO terms included RNA binding and RNA processing.

This consistent elevation of enrichment of functional categories for genes that are more abundant after an RNAlater treatment suggests that this treatment may be altering the physiology of the tissue.

### Genomic Characters Contributing to Differential Expression

We identified four characteristics that contribute significantly to differential gene expression between treatments. Mean expression across samples, GC content, exon number, and homopolymer repeat presence/absence were significant, or nearly significant, terms in the model (Table 2, Figure 3). GC content exhibits the most substantial regression coefficient. The coefficient for GC content is negative, suggesting that genes with higher GC content have a higher relative expression in liquid nitrogen than RNAlater. Mean expression, exon number, and homopolymer repeat presence/absence were significant, or nearly so, such that they exhibited a positive relationship with genes showing higher expression values in RNAlater (i.e., greater mean expression, more exons, having a homopolymer repeat are all related to higher expression in RNAlater). The small regression coefficients of these variable imply, however, that these factors have negligible impacts on differential gene expression observed between preservation methods.

**Table 2:** Terms in the linear model that explain differences in expression between RNAlater store and liquid nitrogen freezing and −80°C storage.

| Term | Sum Sq | Df | F-value | Estimate<br>(SE) | P-value |
| --- | --- | --- | --- | --- | --- |
| Mean Expression | 8 | 1 | 3.4682 | 3.893e-06<br>(1.642e-06) | 0.06258 |
| GC Proportion | 76 | 1 | 31.3766 | -1.092<br>(0.2837) | 2.164e-08 |
| Exon Number | 508 | 1 | 209.9133 | 0.01941<br>(1.340e-03) | <2.2e-16 |
| HPR Presence | 10 | 1 | 4.0495 | 0.05196<br>(0.02582) | 0.04420 |

**Figure 3:**
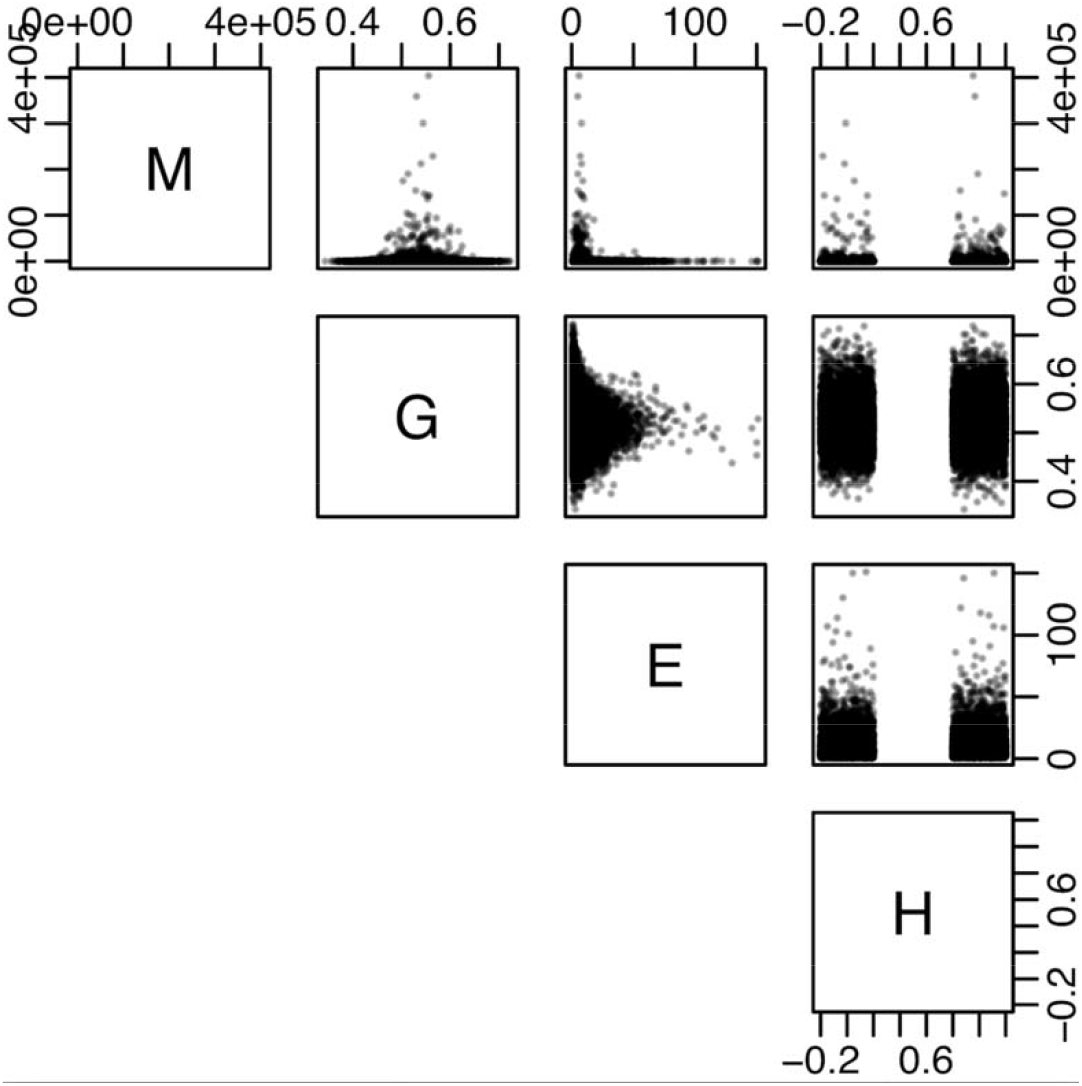
Relationships among the dependent variables retained in the best-fitting generalized linear model. M: mean expression; G: GC content; E: exon number; H: homopolymer repeat presence/absence.

## Discussion

Many sources contribute to variation in observed gene expression. Of these, most researchers are interested in assaying the variation that is due to a biological factor, such as genetic or physiological differences between samples. However, variation due to technical factors, such as noise in hybridization efficacy in microarray studies (Altman 2005) or noise in the number of reads that map to a transcript in RNAseq studies are large sources of variability in observed gene expression, and can substantially influence results (Bryant *et al.* 2011; Marioni *et al.* 2008). For RNA-sequencing studies, the sources of technical variation are still being discovered, but can include many aspects of sample handling prior to actual measurement (McIntyre *et al.* 2011). Previous microarray studies have compared the two sample handling procedures that were tested in our study, and have found no difference downstream, particularly in differential gene expression patterns (Dekairelle *et al.* 2007; Mutter *et al.* 2004). These studies, however, may not apply to the variance profile of RNA-sequencing studies (Romero *et al.* 2012).

Our results suggest that sample handling is an important factor in variation of observed gene expression. While the total percentages of reads mapped were generally similar between the two treatments, the treatments we tested had a significant impact on RNA quality. Our results suggest that preservation in RNAlater, as opposed to flash freezing, non-randomly impacts gene expression values of over 20% of the transcriptome, and our results suggest that shorter genes with higher GC content and lower expression are better preserved in liquid nitrogen. Conversely, our results suggest that genes with high GC content or lower mean expression may not be as well preserved with RNAlater (De Wit *et al.* 2012). The functional enrichment for genes exhibiting significantly higher observed expression in RNAlater than liquid nitrogen indicates that RNAlater may be substantially altering the physiology of the samples during fixation or that RNAlater preserves certain functional categories of genes better than liquid nitrogen. The latter seems more unlikely as it is difficult to hypothesize a mechanism. Further, the converse, does not appear to have extensive enrichment for certain functional categories (i.e., genes that experience presumably worse preservation in RNAlater than liquid nitrogen often do not fall in particular functional categories).

Based on our results, we recommend that researchers use caution when comparing gene expression values derived from RNAseq datasets that may have variable storage conditions. This is especially important with the growth of genomics technologies and accessibility of public data in repositories such as the NCBI Sequence Read Archive. Many entries in these databases do not routinely report metadata such as storage conditions, posing a serious challenge for data utilization. Further, future work could expand on examination of storage in TRIzol (Fisher Scientific, Hampton, NH) as recent work indicates expression patterns might be substantially different from liquid nitrogen (Kono *et al.* 2016). Likewise, various taxonomic groups may be more susceptible to variation in storage conditions because they may exhibit different tissue permeability.

Several caveats are important in interpreting our study. While technical variation from storage condition is the dominant contributor to variation in our study, we acknowledge that biological variation also contributes to our observations. The samples in each storage condition are separate, whole individuals from the same clutch of fish. Fry at 30 days post fertilization are too small to divide tissues equally into preservation treatments and obtain sufficient RNA quantity for RNAseq. Yet, even if a larger tissue sample was cut and divided, one might expect biological variation due to different cell populations. Additionally, juvenile fish tissue may interact with the RNAlater buffer in different ways from other organisms. However, other studies have demonstrated similar effects between RNAlater and flash freezing. For instance, between preservation methods over 5000 differentially regulated genes have been obtained from *Arabidopsis thaliana* tissue (c.f. (Kruse *et al.* 2017)). Though this previous analysis did not assay systematic biases of particular gene attributes to preservation methods, many differentially regulated genes were related to osmotic stress, indicating a strong transcriptional response to RNAlater. Finally, long-term storage temperature is confounded with liquid nitrogen and RNAlater treatments in our study and long-term storage temperature is known to drive RNA integrity (Kono *et al.* 2016) (Gayral *et al.* 2013). Our goal was to replicate typical field experiments, where reliable refrigeration is not available for substantial amounts of time, and RNAlater is used as the predominant preservation method. Despite these caveats, our work demonstrates that differing preservation methods and storage conditions non-randomly impact gene expression, which may bias interpretation of results of RNA sequencing experiments. We look forward to future work that more thoroughly quantifies the impact on interpretation of biological signal derived solely from preservation methods.

## Acknowledgements

We thank the University of Minnesota Genomics Center for their guidance and performing the cDNA library preparations and Illumina HiSeq 2500 sequencing. The authors acknowledge the Minnesota Supercomputing Institute (MSI) at the University of Minnesota for providing resources that contributed to the research results reported within this paper. URL: http://www.msi.umn.edu. Funding was supported by (1R01GM127872-01 to SEM and ACK). CNP was supported by Grand Challenges in Biology Postdoctoral Program at University of Minnesota College of Biological Sciences. Institutional Animal Care and Use Committee at Florida Atlantic University (Protocol #A15-32).

## Data accessibility

All reads are available in NCBI short read archive under accession numbers SRX3446133, SRX3446136, SRX3446135, SRX3446155, SRX3446156, SRS2736519, SRS2736520, SRS2736523, SRS2736524, SRS2736525, and SRS2736526. Scripts to perform all data handling and analysis tasks are available in a GitHub repository at https://github.com/TomJKono/CaveFish_RNAlater

## Supplementary material

**Table S1:** All samples were collected at January 7, 2017 at 10pm EST and were exactly 30-day old fry from the same clutch. RNAlater samples were left on the bench top for 17 days prior to extraction. Liquid N2 samples were flash frozen and stored at −80^□^C prior to extraction. Reported are the treatments (RNALater vs. liquid nitrogen), sample name, extraction date, extraction time, concentration (ng/uL) based on ribogreen, lane the sample was sequenced in and RNA integrity (RIN) scores calculated using RNA bioanalyzer.

| Treatment | Sample | extract_date | extract_time | ng/uL | Lane | RIN |
| --- | --- | --- | --- | --- | --- | --- |
| RNAlater | CHOY-16-R-1 | 1/24/17 | 5:30 PM | 287.53 | 7 | 8.3 |
| RNAlater | CHOY-16-R-2 | 1/24/17 | 5:30 PM | 83.94 | 4 | 8.4 |
| RNAlater | CHOY-16-R-3 | 1/24/17 | 5:30 PM | 39.30 | 8 | 8.8 |
| RNAlater | CHOY-16-R-4 | 1/24/17 | 5:30 PM | 38.71 | 1 | 8.7 |
| RNAlater | CHOY-16-R-5 | 1/24/17 | 5:30 PM | 52.54 | 3 | 8.8 |
| LiquidN2 | CHOY-16-01 | 1/23/17 | 3:30 PM | 264 | 6 | 10.0 |
| LiquidN2 | CHOY-16-04 | 1/24/17 | 1:30 PM | 144.76 | 5 | 9.9 |
| LiquidN2 | CHOY-16-05 | 1/22/17 | 3:30 PM | 69.10 | 8 | 10.0 |
| LiquidN2 | CHOY-16-08 | 1/21/17 | 3:30 PM | 102.10 | 2 | 9.5 |
| LiquidN2 | CHOY-16-11 | 1/19/17 | 12:00 PM | 78.88 | 6 | 10.0 |
| LiquidN2 | CHOY-16-12 | 1/19/17 | 6:00 PM | 67.32 | 5 | 9.6 |

**Figure S1.**
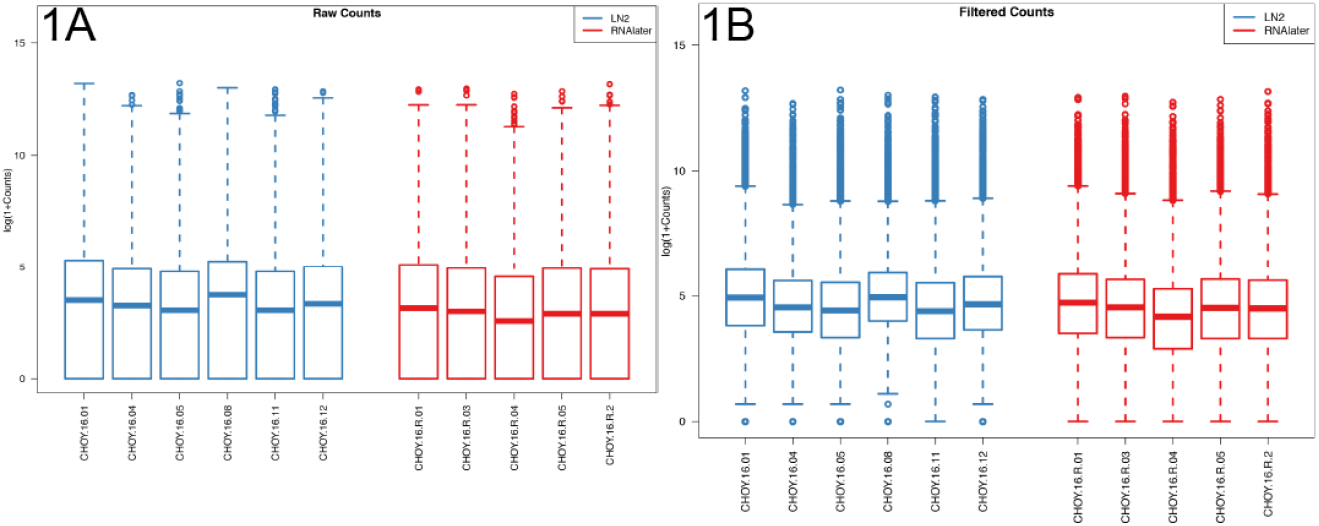
Boxplots depicting normalized counts from DESeq2 for RNALater and liquid nitrogen stored samples. Counts were log transformed (log(1+counts)) for all libraries. A) shows raw counts, and B) shows counts that were filtered for genes with ≤100 counts across all samples.

**Figure S2.**
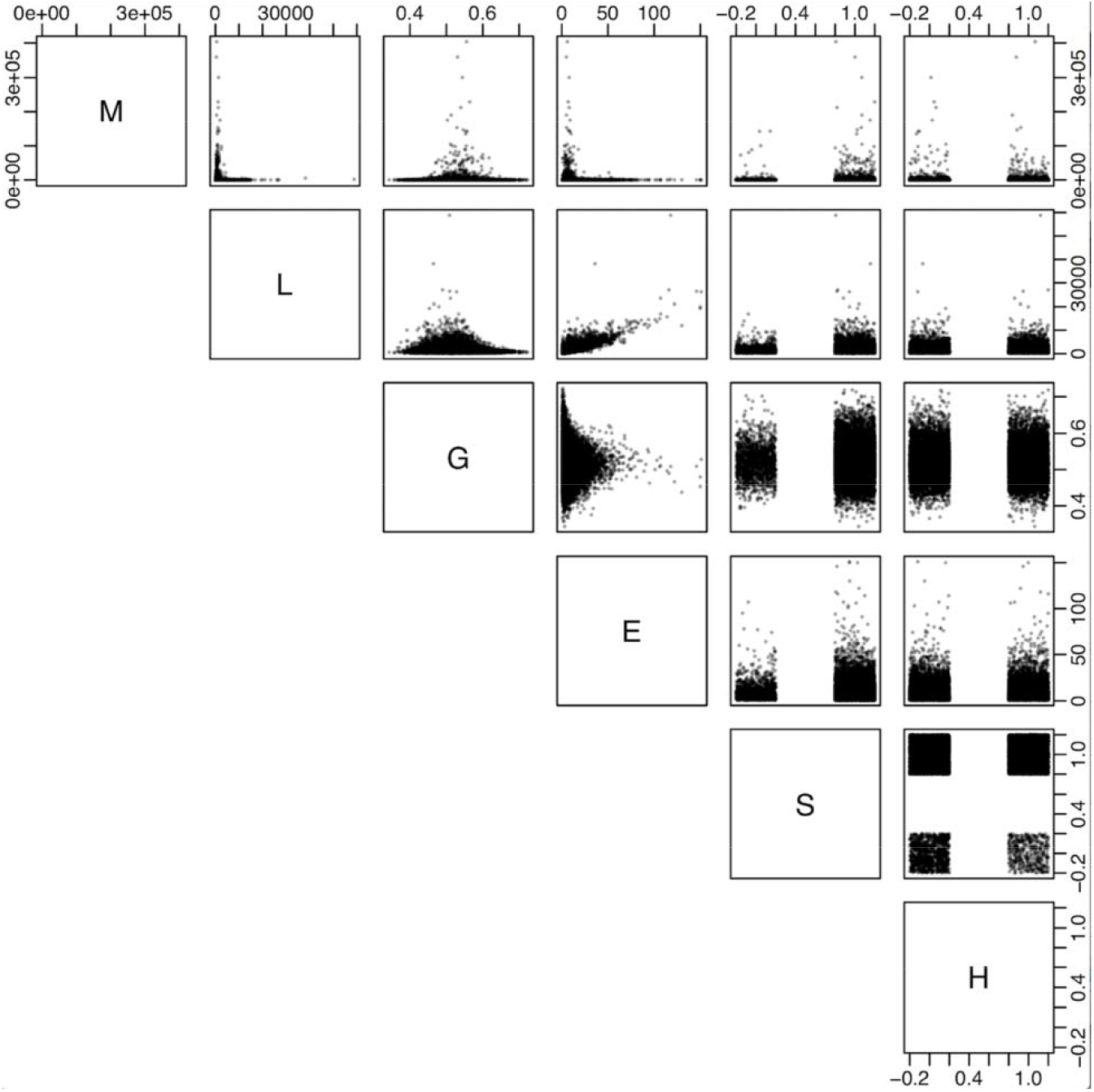
Scatterplot showing the relationships and distributions of all predictors tested in the linear model. M is the mean expression across all samples, L is the annotated gene length, G is the GC content, E is the number of annotated exons, S is simple sequence repeat presence, and H is homopolymer repeat presence. S and H have been jittered to avoid overplotting. Each gene is represented by one point in each scatterplot cell.

**Figure S3.**
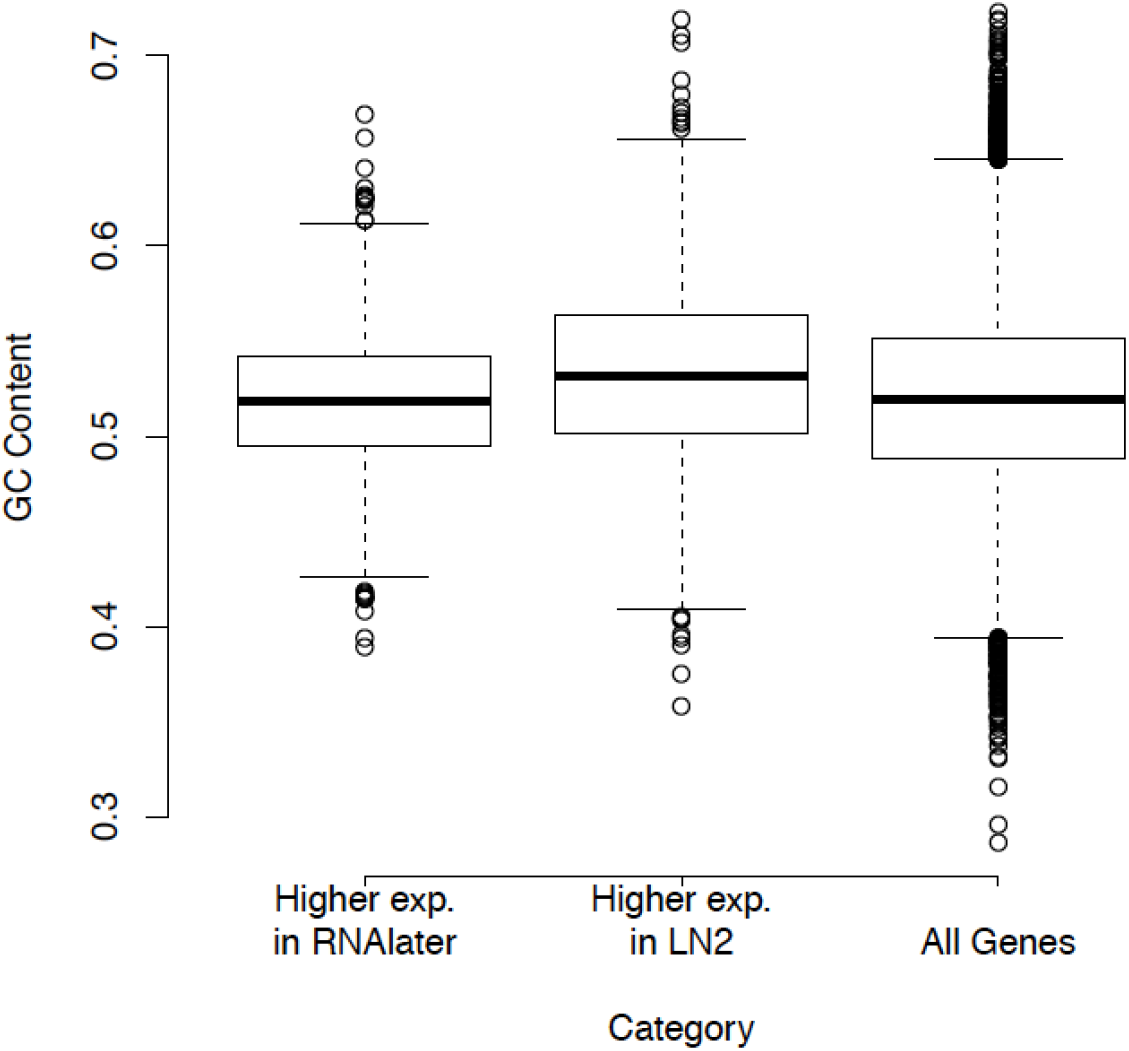
Box plots depicting GC content of genes that were differentially expressed between treatments (e.g. “Higher exp. in”) and all genes that passed the filtering thresholds (e.g. “All Genes”).

